# ABCB1-Mediated Drug Efflux Drives Resistance to VpreB1-Targeted Antibody-Drug Conjugates in B-cell Lymphoblastic Leukemia

**DOI:** 10.64898/2026.08.24.746792

**Authors:** Robin L. Williams, Xiaohong Wang, Jason Ostergaard, Jongseok Kang, Madison Gohman, Luke Lambert, Timothy Singleton, Sarah K. Tasian, Megan Hilgers, Keegan C. Lee, Joseph M. Muretta, Stuart S. Winter, Peter M. Gordon

**Author notes:** Contact information for corresponding author: Peter Gordon, M.D./Ph.D., Division of Pediatric Hematology/Oncology University of Minnesota, 420 Delaware St SE, MMC 366, Minneapolis, MN, 55455. RLW and XW contributed equally to this work. **Conflict of Interest Statement:** The authors declare no potential conflicts of interest.

## Abstract

**Aim:** Although B-cell acute lymphoblastic leukemia (B-ALL) is highly responsive to antigen-directed immunotherapies, treatment resistance remains a major barrier to achieving durable responses in patients. We recently developed a novel VpreB1 (CD179a)-directed antibody-drug conjugate with calicheamicin (VpreB1-ADC) that exploits the restricted expression of VpreB1 within the surrogate light chain in early B cells, including B-ALL. In the present work, we investigated mechanisms of resistance to the VpreB1-ADC.

**Methods:** Mechanisms of resistance were evaluated using a *TCF3::HLF* B-ALL model, assessing target engagement parameters including VpreB1 surface expression and antibody internalization. The role of the multidrug resistance transporter ABCB1 (P-glycoprotein) was evaluated via pharmacologic inhibition, using tariquidar and zosuquidar, and enforced overexpression across multiple B-ALL cell lines. Sensitivity to alternative non-ABCB1 substrate payloads exatecan and PNU-159682 was also assessed.

**Results:** Resistant *TCF3: HLF* cells retained VpreB1 expression and efficient antibody internalization. Instead, resistance was driven by elevated ABCB1 expression and activity. ABCB1 inhibition with tariquidar or zosuquidar restored VpreB1-ADC sensitivity. Conversely, enforced ABCB1 overexpression conferred ADC resistance, which was reversed by ABCB1 inhibition. Cells with high ABCB1 activity remained fully sensitive to alternative payloads, including exatecan and PNU-159682, which are not ABCB1 substrates.

**Discussion:** ABCB1-mediated drug efflux drives intrinsic resistance to calicheamicin-conjugated ADCs in B-ALL. Combining ADCs with ABCB1 inhibitors or selecting payloads non-susceptible to ABCB1 efflux offer viable strategies to overcome resistance and optimize future ADC therapies.

## Introduction

B-lineage acute lymphoblastic leukemia/lymphoma (B-ALL/B-LL or B-LBL), the most common malignancy in children and young adults, is highly curable with intensive chemotherapy, but outcomes remain poor in high-risk or relapsed disease and are limited by substantial treatment-related toxicity[1,2]. These limitations have driven the development of targeted immunotherapies, including antibodies, antibody-drug conjugates (ADCs), bispecific T cell engagers (BiTEs), and chimeric antigen receptor T cells (CAR-T), which have demonstrated strong clinical efficacy and underscore the immunoresponsive nature of B-ALL[3]. However, important challenges persist, including treatment-related toxicities and emerging resistance mechanisms that constrain the durability and breadth of responses[4].

We recently developed an ADC targeting VpreB1 (CD179a), a component of the surrogate light chain (SLC) which is transiently expressed during early B-cell development[5]. Together with IGLL1 (λ5, CD179b), VpreB1 mediates pre-B-cell receptor (pre-BCR) assembly and signaling, which drives early B-cell expansion before SLC expression is extinguished. Notably, VpreB1 is broadly expressed across B-ALL subtypes and persists in minimal residual disease, while being absent from mature B cells, supporting its potential as a selective therapeutic target[6].

Consistent with this rationale, the unconjugated VpreB1 antibody showed minimal activity, whereas the VpreB1 antibody conjugated to calicheamicin induced potent, dose-dependent cytotoxicity across multiple B-ALL cell lines and patient-derived xenograft (PDX) models comprised of various genetic backgrounds. *In vivo*, treatment reduced leukemic burden, prolonged survival, and induced durable remissions, including long-term cures in a subset of animals.

However, responses were not uniform. HAL-01 and ALL1807 cells (immortalized for *in vitro* studies from an *in vivo* PDX model[7,8]), which harbor the rare t(17;19) translocation generating the *TCF3*::*HLF* fusion, exhibited significant resistance. This aligns with the aggressive biology of *TCF3*::*HLF* B-ALL, which is associated with chemotherapy resistance and very poor prognosis, potentially mediated by multidrug efflux transporters and anti-apoptotic signaling[9–12]. These findings highlight both a limitation of the VpreB1-ADC and an opportunity to define resistance mechanisms. Accordingly, we hypothesized that *TCF3*::*HLF* B-ALL models could reveal determinants of VpreB1-ADC response, with broader implications for ADC resistance.

Herein, we identify ABCB1-mediated drug efflux as a mechanism of resistance to VpreB1-ADC in B-ALL and demonstrate that this resistance can be reversed with pharmacologic ABCB1 inhibition or potentially circumvented using alternative ADC payloads that are not ABCB1 substrates. More broadly, our findings implicate multidrug resistance (MDR) transporters as important determinants of ADC response in B-ALL and support the development of combinatorial or payload-optimized strategies to overcome resistance in high-risk disease.

## Materials and Methods

### Cell culture

Commercially available leukemia cell lines were from the American Type Culture Collection (REH) or the German Collection of Microorganisms and Cell Cultures GmbH (HAL-01, NALM-6, and SEM) and cultured in RPMI media supplemented with fetal bovine serum (FBS; Seradigm) and penicillin-streptomycin. The ALL1807 PDX model was established in NSG mice and immortalized for *in vitro* cell culture as previously described[7,8].

### VpreB1-ADC synthesis

As previously described, a humanized monoclonal anti-VpreB1 antibody (IgG1) was produced by Genscript and conjugated via lysine linkage using an acid-labile acetyl butyrate linker to N-acetyl γ-calicheamicin (Creative Biolabs), yielding a drug:antibody ratio of 4.97, aggregation of 1.63%, and endotoxin <1 EU/mg[5].

### Drugs, antibodies, and reagents

Tariquidar, zosuquidar, vincristine, doxorubicin, cytarabine, PNU-159682, exatecan, and calcein-AM were purchased from MedChemExpress. A phycoerythrin (PE)-conjugated anti-human ABCB1 (CD243/P-gp) mouse monoclonal antibody (clone UIC2; BioLegend, #348605) and a mouse IgG2a, κ isotype control antibody (clone MOPC-173; BioLegend, #400213) were obtained from BioLegend. Flow cytometry analyses of primary B-ALL samples used the following antibodies from BD Pharmigen: PE mouse anti-human P-glycoprotein (CD243/ABCB1), BV711 mouse anti-human CD338 (ABCG2/BCRP), FITC mouse anti-human MRP1(ABCC1).

### Intracellular anti-VpreB1 antibody uptake

Experiments were performed as previously described[5]. In brief, leukemia cells were labeled with both a fluorescent (AF647) and biotin conjugated anti-VpreB1 antibody. Internalization was then either blocked by incubation on ice or allowed to proceed for 1 hour at 37 °C. Cells were then labeled with NeutrAvidin-FITC (Thermo Fisher #A2662) and assessed by flow cytometry. FITC and AF647 (APC) fluorescence indicate cell-surface anti-VpreB1and total anti-VpreB1 (cell-surface plus intracellular), respectively.

### Proliferation and apoptosis analyses

Leukemia cells were cultured in 96-well plates and viability assessed with the CellTiter-Glo Luminescent Cell Viability Assay (Promega) and a Tecan Infinite M200 Pro plate reader. All experiments were performed with at least 3 wells per condition. To assess apoptosis and cell death, leukemia cells were stained with annexin-V antibody (eBioscience), a fixable viability dye (ThermoFisher Scientific), and analyzed by flow cytometry using a BD Accuri C6 or Symphony instrument.

### Calcein-acetoxymethyl ester (calcein-AM) staining

Leukemia cells (2×10⁵/mL) were treated *in vitro* with DMSO, tariquidar (1 µM), or zosuquidar (1 µM) for 5 hours. Calcein-AM was then added to a final concentration of 25 nM, and cells were incubated for an additional 24 hours.

Cells were harvested, washed, and stained with eBioscience™ Fixable Viability Dye eFluor™ 660 for 5 min. Calcein fluorescence in viable cells was analyzed by flow cytometry after washing and resuspension in FACS buffer.

### Lentiviral production and transduction of leukemia cells

A lentiviral vector and particles (>10^8^ TU/mL) expressing humanized ABCB1 under control of the SFFV promoter was purchased from VectorBuilder (VectorBuilder Inc, USA). The vector also expressed enhanced green fluorescent protein (EGFP) and puromycin to facilitate cell selection. Vectors were verified by sequencing. Leukemia cells adhered to retronectin-coated tissue culture plates were transduced with virus particles (MOI 10) for 48 hours. Transduced cells were enriched by puromycin selection and then sorted by flow cytometry on a BD FACS Aria II instrument for EGFP expression.

### Statistical analysis

Data are presented as mean ± SD. All experiments were performed at least three times with representative data of one experiment presented. For statistical analysis, Student’s t-tests were used for two-group comparisons, while one-way ANOVA was employed for multiple groups. Curve fitting (four-parameter logistic non-linear regression) and statistical analyses were performed using GraphPad Prism 11 (GraphPad Software, La Jolla, CA). Significance levels are denoted in figure legends (*ns*, not significant; *, *P* < 0.05; **, *P* < 0.01; ***, *P* < 0.001; ****, *P* < 0.0001), with *P* < 0.05 considered statistically significant.

## Results

Consistent with our prior findings, HAL-01 cells were completely resistant to the VpreB1-ADC, whereas ALL1807 cells exhibited relative resistance compared to NALM-6 leukemia cells (Figure 1A)[5]. Moreover, despite their relative resistance, both cell lines internalized the VpreB1 antibody (Figure 1B), indicating that neither impaired antigen expression nor defective internalization was likely to account for the observed treatment resistance.

**Figure 1:**
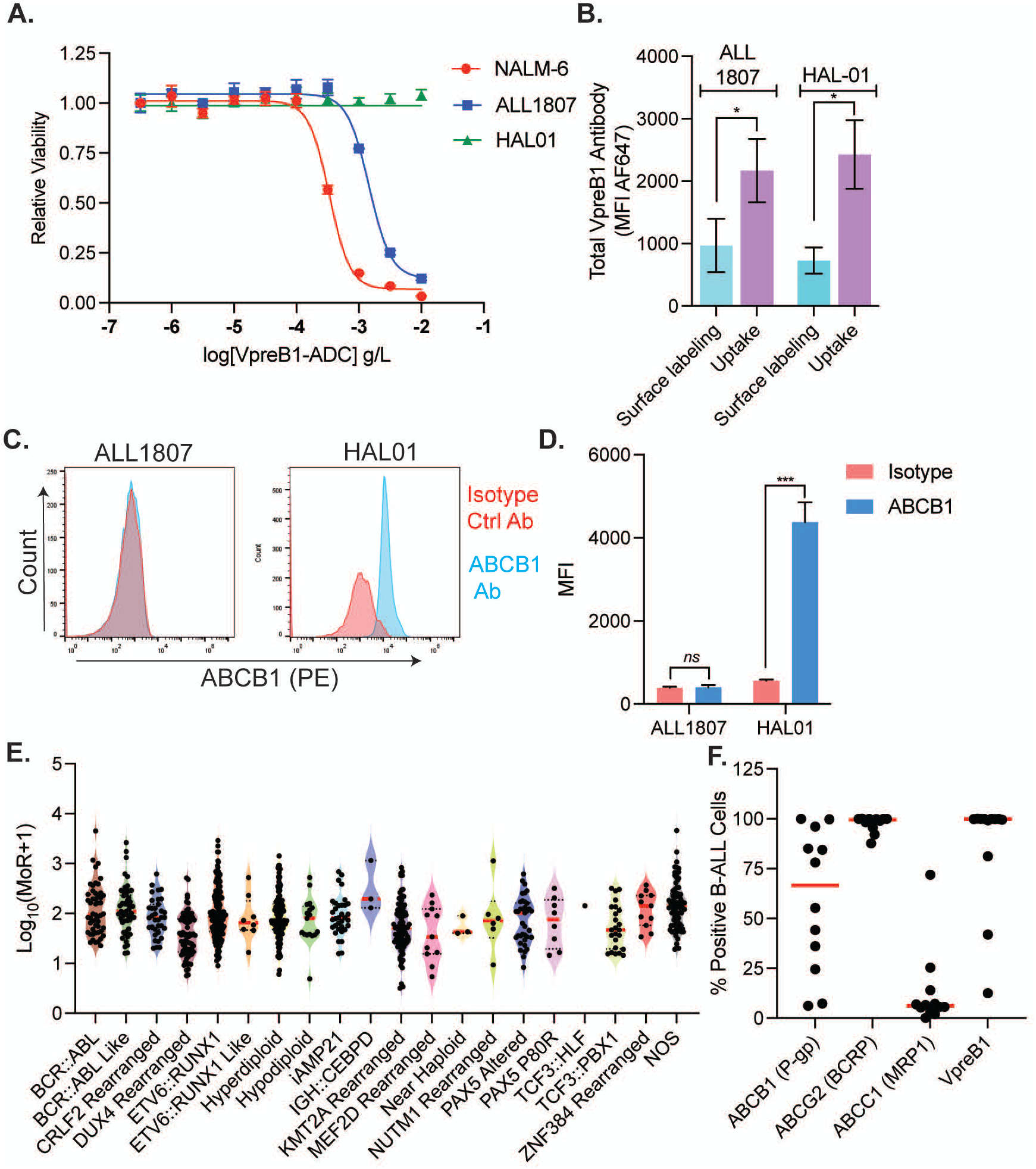
ABCB1 expression in B-ALL is associated with VpreB1-ADC resistance. **A**. *In vitro* VpreB1-ADC dose-response curves for NALM-6, ALL1807, and HAL-01 leukemia cell lines. Leukemia cell viability was assessed after 42 hours of ADC treatment using the CellTiter-Glo Luminescent Cell Viability Assay, which quantifies ATP as a measure of metabolically active cells. Error bars represent the mean ± SD of three technical replicates. **B.** Internalization of cell surface VpreB1 protein on HAL-01 and ALL1807 cells was measured by labeling the cells with a fluorescent (AF647) anti-VpreB1 antibody. Internalization was then either blocked by incubation on ice or allowed to proceed for 1 hour at 37 °C. Error bars represent the mean ± SD and *P*, *, <0.05 by t-test. **C-D**. ALL1807 and HAL-01 cells were stained with either ABCB1 or isotype control antibody and assessed by flow cytometry. Representative flow cytometry histograms are shown in (**C**) and median fluorescent intensity (MFI) quantitation in (**D)**. **E**. Assessment of *ABCB1* mRNA expression across molecular subtypes of >800 primary pediatric and young adult B-ALL samples using RNA-seq data from the St. Jude Cloud database (https://www.stjude.cloud). **F.** Expression of VpreB1 and the drug efflux transporters ABCB1 (P-gp), ABCG2 (BCRP), and ABCC1 (MRP1) was assessed by flow cytometry on diagnostic bone marrow samples from consecutive newly diagnosed pediatric B-ALL patients.

Given prior reports implicating the MDR transporter P-glycoprotein (P-gp; ABCB1) in the known chemoresistance of *TCF3*::*HLF* B-ALL, we next evaluated ABCB1 expression[11]. HAL-01, but not ALL1807 cells, exhibited high ABCB1 protein levels (Figure 1C-D). To assess ABCB1 expression in B-ALL more broadly, we analyzed *ABCB1* mRNA expression levels in >800 primary B-ALL samples from pediatric and young adult patients in the publicly available St. Jude Cloud database[13]. As shown in Figure 1E, *ABCB1* expression was generally intermediate across most samples, but a subset exhibited more elevated levels. Notably, only a single sample harbored an *HLF* rearrangement, precluding confirmation of the prior report linking *TCF3*::*HLF* to elevated *ABCB1* mRNA expression. To account for potential discordance between RNA and protein expression, we next measured ABCB1 (P-gp) protein expression by flow cytometry on diagnostic bone marrow samples from 12 children, adolescents, or young adults with newly diagnosed B-ALL. We expanded this evaluation to include two additional ATP-binding cassette transporters known to mediate MDR, ABCG2 (BCRP) and ABCC1 (MRP1). All samples exhibited VpreB1 expression, predominantly at high levels (range 12.5–100%; median 99.8%). ABCB1 was expressed at variable levels across all patient samples (range 6.17–99.9%; median 66.7%). While ABCG2 was strongly expressed in most samples (range 87.7–99.9%; median 99.6%), ABCC1 expression remained predominantly low (range 0.18–71.9%; median 6.2%) (Figure 1F and Supplemental Table 1). Collectively, these data suggest that ABCB1 expression may mediate resistance to VpreB1-ADCs in HAL-01 cells and potentially across a subset of B-ALL cases.

We next sought to obtain functional evidence supporting a role for ABCB1 in ADC resistance. Accordingly, we evaluated the effect of the third-generation ABCB1 inhibitor tariquidar on drug efflux and VpreB1-ADC sensitivity[14–16]. Prior to testing VpreB1-ADC, we first used calcein-AM, a non-fluorescent, membrane-permeant ABCB1 substrate that can be effluxed by ABCB1 prior to intracellular esterase-mediated conversion to the fluorescent, membrane-impermeant calcein. Intracellular fluorescence thus served as an inverse readout of ABCB1 activity.

Tariquidar was non-toxic to leukemia cells under the conditions tested (Figure 2A). Consistent with the expression data, HAL-01, but not ALL1807, exhibited efficient efflux of calcein-AM, confirming elevated functional ABCB1 activity (Figure 2B-C). More importantly, tariquidar completely sensitized HAL-01, but not ALL1807, cells to VpreB1-ADC, reducing viability from >90% to <5% (Figure 2D-G). Moreover, a transient overnight exposure to tariquidar, followed by drug washout for up to 96 hours prior to VpreB1-ADC administration, was still sufficient to sensitize HAL-01 cells, indicating a durable impact of ABCB1 inhibition on ADC sensitivity (Figure 2H).

**Figure 2:**
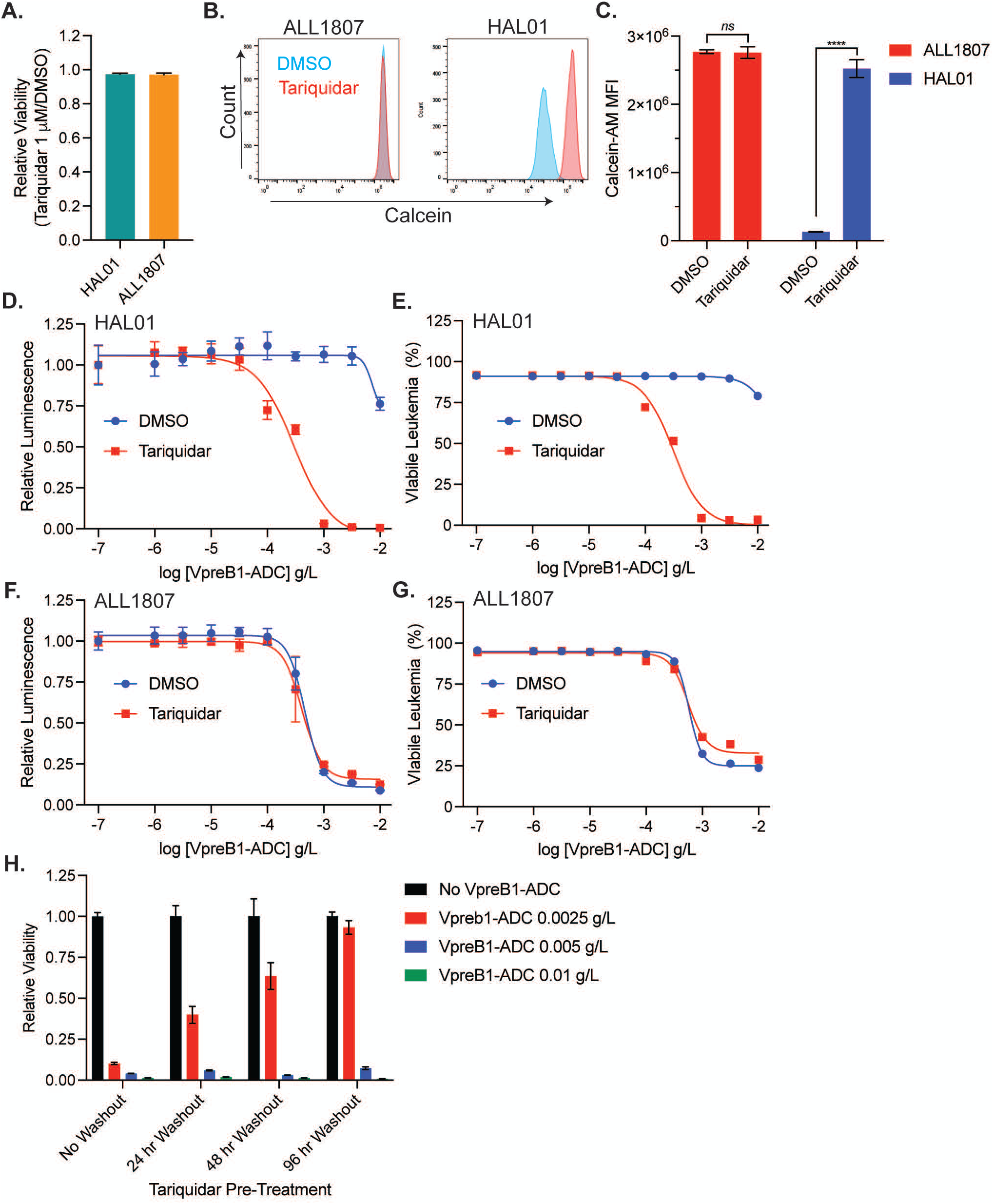
Pharmacologic inhibition of ABCB1 restores VpreB1-ADC sensitivity in resistant TCF3::HLF B-ALL cells. **A.** HAL-01 and ALL1807 cells were treated with tariquidar 1 μM or DMSO. Viability was assessed after 48 hours of drug treatment using the CellTiter-Glo Luminescent Cell Viability Assay. Error bars represent the mean ± SD of three technical replicates. **B-C.** Representative flow cytometry histograms (**B**) and median fluorescent intensity (MFI) quantitation (**C**) of intracellular calcein fluorescence in ALL1807 (left) and HAL-01 (right) cells treated with either DMSO (blue) or tariquidar (red). Error bars represent the mean ± SD of three technical replicates and *P*: *ns*, not significant, ****, <0.0001. **D-G**. VpreB1-ADC dose-response curves for HAL-01 (**D, E**) and ALL1807 (**F, G**) leukemia cells in the presence of tariquidar 1 μM or DMSO control. Leukemia cell viability was measured after 48 hours using the CellTiter-Glo Assay (**D, F**) or annexin-V and viability dye staining followed by flow cytometry (**E, G**). Error bars represent mean ± SD of three technical replicates. **H**. HAL-01 cells were pretreated with tariquidar 1 µM for 12 hours, washed three times with PBS, and then cultured in fresh media for the indicated duration prior to VpreB1-ADC exposure. After 48 hours of drug treatment, leukemia cell viability was assessed using the CellTiter-Glo assay.

To validate a causal role for ABCB1 in VpreB1-ADC resistance, we overexpressed ABCB1 in ALL1807 cells and in additional VpreB1-ADC-sensitive B-ALL cell lines with low endogenous ABCB1 expression, NALM-6 and REH (Supplemental Figure 1). Consistent with our observations in HAL-01 cells, tariquidar increased calcein retention in ABCB1-overexpressing cells, confirming functional inhibition of ABCB1-mediated efflux (Supplemental Figure 2).

Importantly, ABCB1 overexpression markedly reduced sensitivity to VpreB1-ADC across all tested cell lines. However, this resistance was fully reversed by co-treatment with tariquidar (Figure 3), further supporting a direct role for ABCB1-mediated efflux in limiting VpreB1-ADC cytotoxicity To better assess the sensitivity of ABCB1 to pharmacologic inhibition in leukemia cells, we generated tariquidar dose-response curves in HAL-01 cells and in ABCB1-overexpressing leukemia cell lines in the presence of vehicle control or a cytotoxic concentration of VpreB1-ADC (Figure 4A-D). From these dose-response curves, we calculated that the tariquidar IC_50_ ranged from approximately 10 to 100 nM across the different leukemia cell lines (Figure 4E). Further supporting these findings, zosuquidar, a distinct third-generation ABCB1 inhibitor, similarly enhanced the sensitivity of both HAL-01 cells and ABCB1-overexpressing leukemia cell lines to VpreB1-ADC (Supplemental Figure 3), with an IC_50_ ranging from 20 to 500 nM (Supplemental Figure 4).

**Figure 3:**
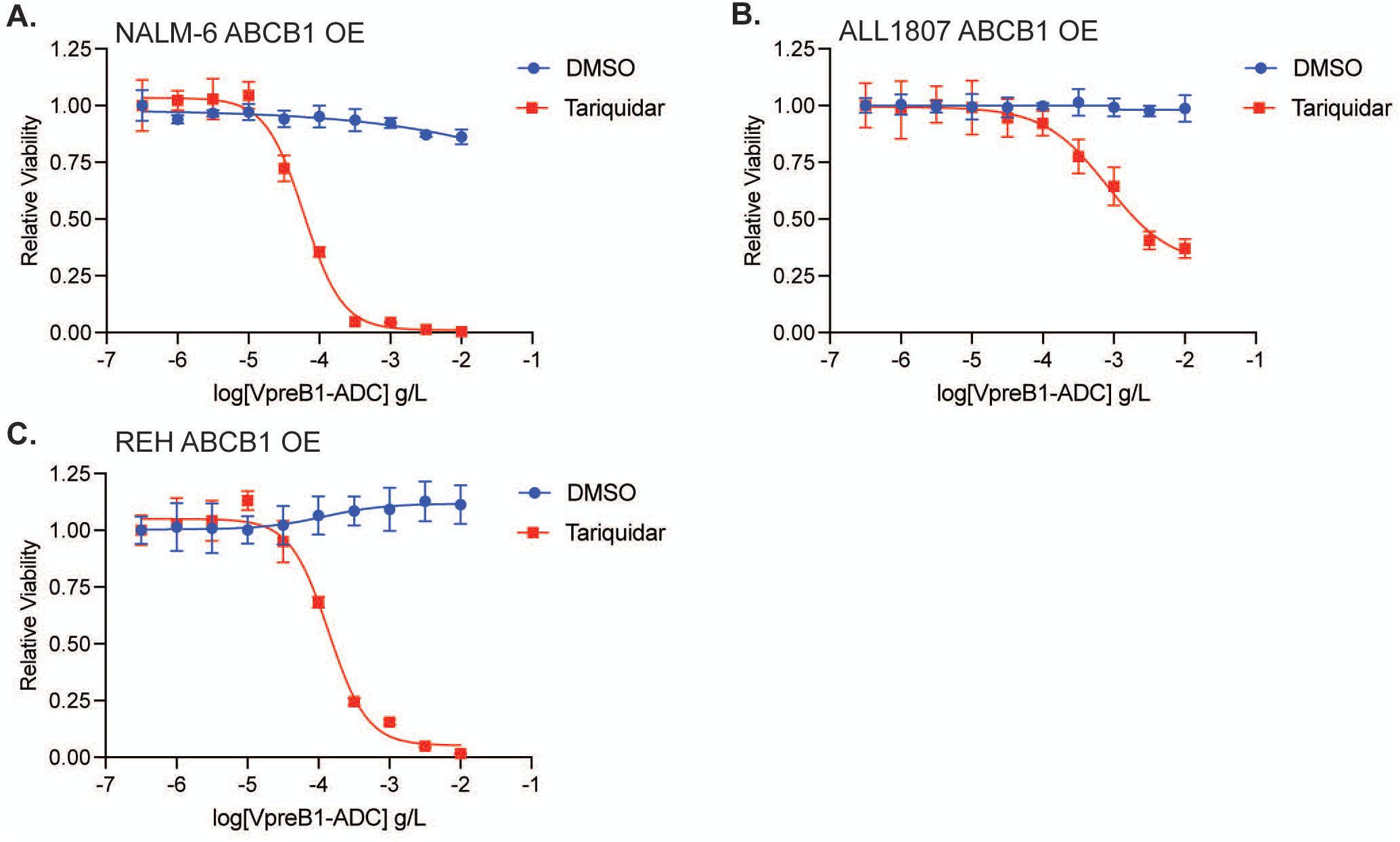
ABCB1 overexpression confers VpreB1-ADC resistance that is reversible with tariquidar treatment. **A-C.** VpreB1-ADC dose-response curves for NALM-6 (**A**), ALL1807 (**B**), and REH (**C**) leukemia cells engineered to overexpress ABCB1 and treated with either tariquidar 1 μM or vehicle control. Leukemia cell viability was measured after 48 hours using the CellTiter-Glo Assay. Error bars represent mean ± SD of three technical replicates.

**Figure 4:**
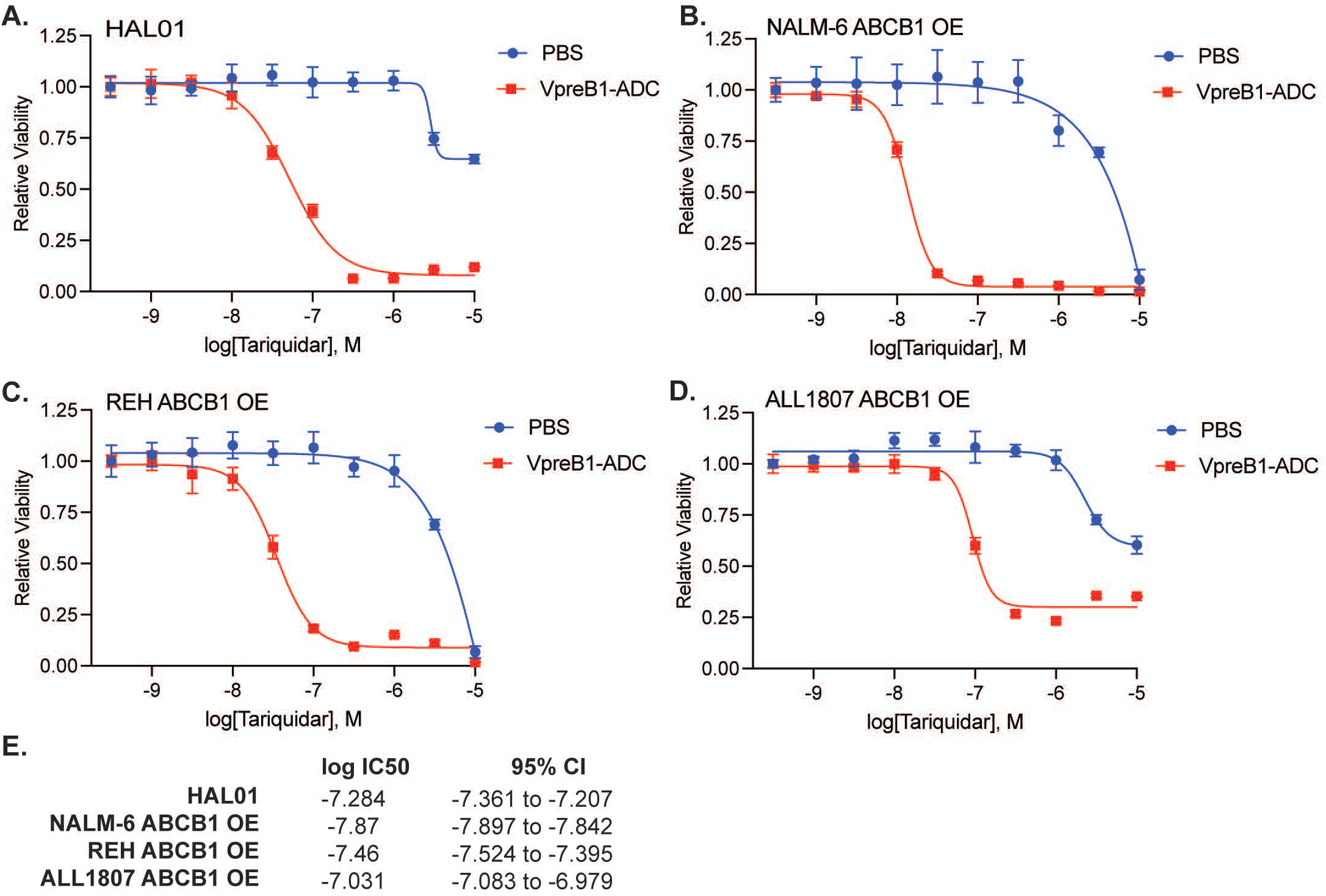
Tariquidar IC₅₀ determination. **A-D**. Tariquidar dose-response curves for HAL-01 (**A**), NALM-6 ABCB1-overexpressing (**B**), REH ABCB1-overexpressing (**C**), and ALL1807 ABCB1-overexpressing (**D**) leukemia cell lines in the presence of VpreB1-ADC 0.001 g/L or vehicle control. Leukemia cell viability was measured after 48 hours using the CellTiter-Glo Assay. Error bars represent mean ± SD of three technical replicates. **E**. Tariquidar IC₅₀ values calculated from the dose-response curves.

To determine whether the observed effects of ABCB1 overexpression were specific to VpreB1-ADC or reflected a broader, substrate-dependent drug resistance phenotype, we evaluated the response of ABCB1-overexpressing leukemia cells to additional chemotherapeutic agents that are either established ABCB1 substrates (vincristine and doxorubicin) or not (cytarabine)[17]. As predicted, ABCB1 overexpression conferred marked resistance to vincristine and doxorubicin, but had less of an impact on cytarabine sensitivity (Supplemental Figure 5). Notably, this resistance to vincristine and doxorubicin was reversed by co-treatment with either tariquidar or zosuquidar (Supplemental Figure 6), further supporting the conclusion that ABCB1-mediated drug efflux is responsible for the observed resistance phenotype. Finally, we evaluated loncastuximab tesirine, a CD19-targeted ADC approved by the FDA for adults with relapsed/refractory diffuse large B-cell lymphoma, which delivers the DNA alkylating payload tesirine (SG3199), a weak-to-moderate ABCB1 substrate[18,19]. Consistent with our VpreB1-ADC findings, tariquidar significantly enhanced loncastuximab cytotoxicity in HAL-01 and ABCB1-overexpressing leukemia cell lines (Supplemental Figure 7), consistent with ABCB1-mediated efflux of the tesirine payload. However, the magnitude of sensitization was less pronounced in HAL-01 and ALL1807 cells overexpressing ABCB1, suggesting that additional determinants beyond ABCB1 may influence tesirine sensitivity in these cell lines. Together, these results indicate that ABCB1 expression confers a broader, substrate-dependent resistance phenotype in B-ALL that extends beyond VpreB1-ADC to other ADCs and conventional chemotherapeutics, while remaining at least partially reversible with pharmacologic inhibition of ABCB1.

Finally, as an alternative to combining the VpreB1-ADC with ABCB1 inhibition, we next evaluated additional cytotoxic agents that have been successfully incorporated as ADC payloads but are not considered ABCB1 substrates, specifically the free drugs exatecan [20,21] and PNU-159682[22,23]. In contrast to the calicheamicin-conjugated VpreB1-ADC, HAL-01 and ABCB1-overexpressing leukemia cell lines exhibited marked sensitivity to both exatecan and PNU-159682, and this sensitivity was not further enhanced by co-treatment with tariquidar (Figures 5-6), consistent with these agents largely evading ABCB1-mediated drug efflux.

**Figure 5:**
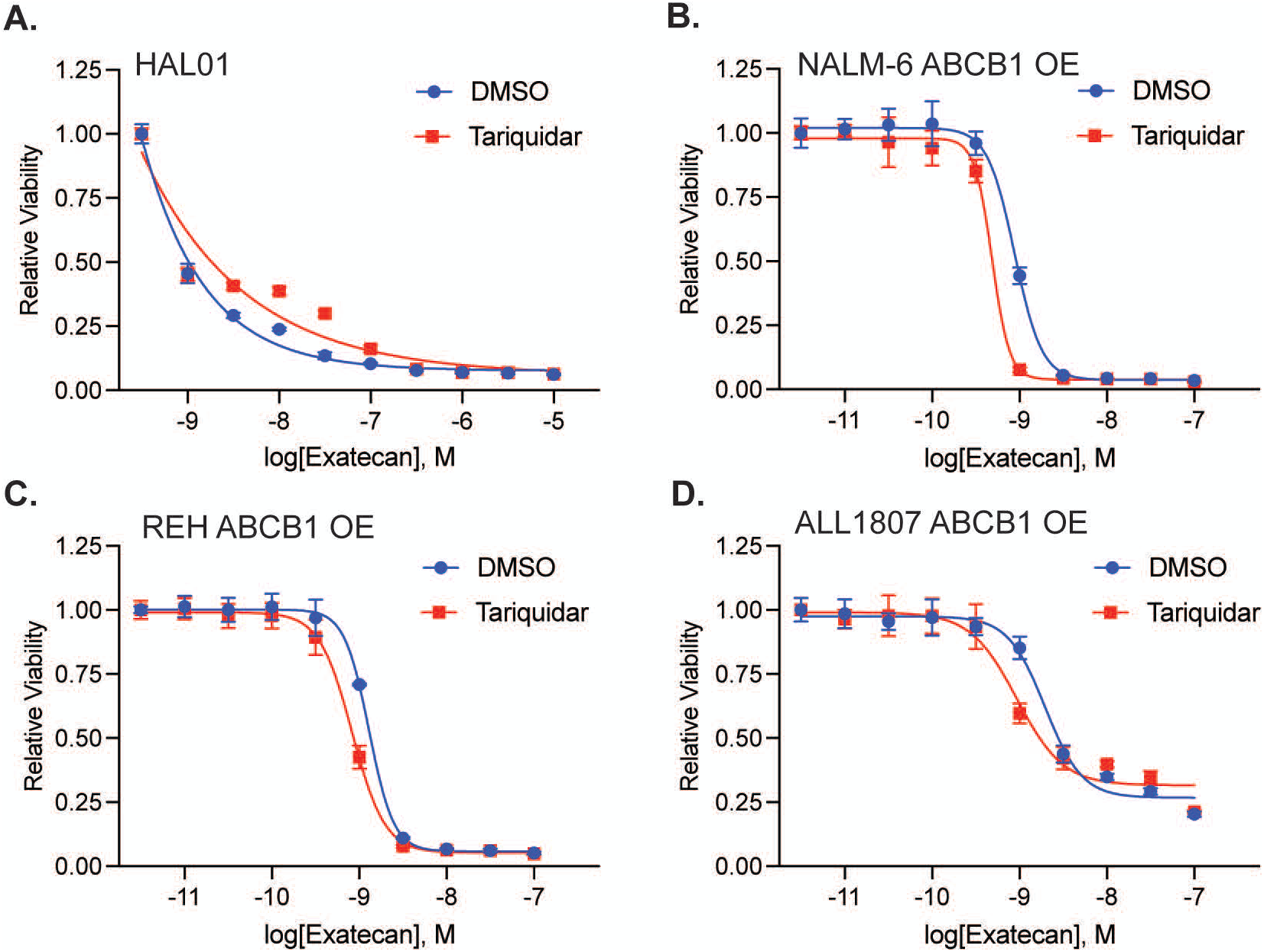
Exatecan retains cytotoxic activity in B-ALL cells despite ABCB1 expression. **A-D.** Exatecan dose-response curves for HAL-01 (**A**), NALM-6 ABCB1-overexpressing (**B**), REH ABCB1-overexpressing (**C**), and ALL1807 ABCB1-overexpressing (**D**) leukemia cell lines in the presence of tariquidar 1 μM or DMSO control. Leukemia cell viability was measured after 48 hours using the CellTiter-Glo Assay. Error bars represent mean ± SD of three technical replicates.

**Figure 6:**
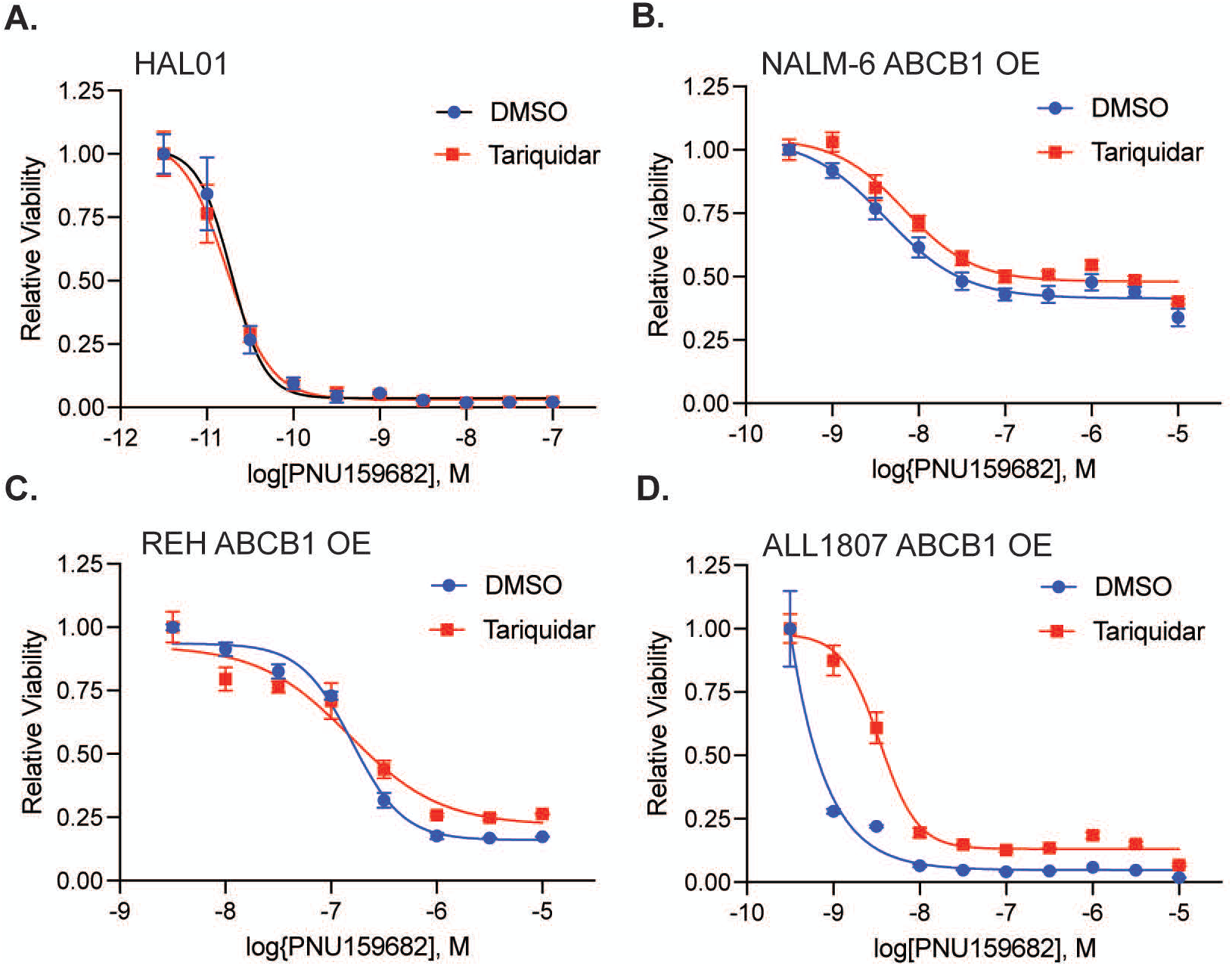
PNU159682 retains cytotoxic activity in B-ALL cells despite ABCB1 expression. **A-D**. PNU159682 dose-response curves for HAL-01 (**A**), NALM-6 ABCB1-overexpressing (**B**), REH ABCB1-overexpressing (**C**), and ALL1807 ABCB1-overexpressing (**D**) leukemia cell lines in the presence of tariquidar 1 μM or DMSO control. Leukemia cell viability was measured after 48 hours using the CellTiter-Glo Assay. Error bars represent mean ± SD of three technical replicates.

## Discussion

In the present study, we sought to define mechanisms of B-ALL resistance to a VpreB1-directed ADC (VpreB1-ADC). Consistent with our prior work, insensitivity to our novel VpreB1-ADC was observed in select cell lines, most notably two harboring *TCF3*::*HLF* fusions [5]. Importantly, we found that VpreB1-ADC resistance in HAL-01, but not ALL1807 cells, was driven by active drug efflux mediated by the MDR transporter ABCB1 (P-glycoprotein). Pharmacologic inhibition of ABCB1 fully restored VpreB1-ADC sensitivity in HAL-01 and other ALL cell lines engineered to overexpress ABCB1, establishing a therapeutically actionable mechanism of resistance.

ABCB1-mediated drug efflux has long been implicated in chemotherapy resistance in acute leukemias, including B-ALL[24]. Reported ABCB1 positivity rates in B-ALL vary widely across studies, from <10% to >50%, likely due to small cohort sizes, *de novo* versus relapsed patient populations, differences in detection methods (RNA vs protein vs functional assays), antibody selection, and thresholds for positivity, making the true incidence difficult to define[25–32]. In our study, flow cytometric analysis of diagnostic bone marrow samples from newly diagnosed pediatric and AYA B-ALL patients demonstrated detectable ABCB1 expression in all cases, although levels varied substantially between samples. In contrast, ABCG2 expression was more uniformly high across most specimens, whereas ABCC1 expression was generally lower and more heterogeneous, highlighting the diversity of MDR transporter expression patterns in primary B-ALL.

While some studies link increased ABCB1 expression to higher relapse risk and inferior survival of patients with B-ALL[26–28], others show no prognostic significance[25,31,32]. This inconsistency, coupled with a lack of methodological standardization for defining positivity, precludes firm conclusions about the clinical relevance of ABCB1 in B-ALL. Nonetheless, the collective data, together with our analyses of ABCB1 expression in >800 B-ALL patient samples from the St. Jude database and our flow cytometric characterization of 12 primary B-ALL specimens, indicate that a meaningful subset of B-ALL patients express ABCB1. Moreover, assessment of ABCB1 in diagnostic specimens likely underestimates its importance in B-ALL, as protein expression can be upregulated following chemotherapy and through interactions with the bone marrow microenvironment[33,34]. Finally, ABCB1 is just one of several MDR transporters with distinct yet often overlapping substrate and inhibition specificities, suggesting that efflux-mediated resistance in B-ALL is multifactorial and not fully captured by evaluating ABCB1 alone[35]. Consistent with this concept, we observed high ABCG2 expression and variable ABCC1 expression across primary B-ALL samples, raising the possibility that these transporters may also contribute to resistance to specific chemotherapeutic agents or ADC payloads depending on their substrate profiles.

Our findings extend this paradigm to ADCs, demonstrating that even when antigen targeting and internalization are intact, intracellular payload delivery can be effectively neutralized by efflux transporters. This is particularly relevant for ADCs utilizing payloads such as calicheamicin, which are known substrates of ABCB1. These data strongly support a model in which intracellular payload retention, in addition to antibody binding and uptake, represents a critical determinant of ADC efficacy.

The clinical targeting of ABCB1-mediated resistance in cancer therapy has been explored extensively, although early efforts to therapeutically inhibit MDR transporters were largely unsuccessful. First- and second-generation ABCB1 inhibitors were limited by poor specificity, dose-limiting toxicities, and adverse pharmacokinetic interactions[36–38]. In contrast, third-generation inhibitors such as tariquidar and zosuquidar, used in this study, demonstrate improved potency, selectivity, and tolerability, with minimal effects on cytochrome P450-mediated drug metabolism[14,17,39]. These advances have renewed interest in targeting ABCB1 and, more broadly, MDR proteins as a therapeutic strategy. Supporting this approach, studies in acute myeloid leukemia (AML) have implicated MDR transporters in resistance to gemtuzumab ozogamicin, a CD33-directed ADC. Notably, a recent clinical trial demonstrated improved outcomes in ABCB1-positive patients treated with gemtuzumab ozogamicin, which employs the same calicheamicin payload used in our construct, when combined with zosuquidar, relative to ABCB1-negative patients.[39–42]. Together with our findings, these data suggest that combining ADCs with MDR inhibitors may represent a rational and broadly applicable strategy to overcome resistance in leukemia.

Our findings also suggest alternative strategies to overcome ABCB1-mediated resistance beyond pharmacologic inhibition. Specifically, we show that leukemia cells resistant to the calicheamicin-based VpreB1-ADC remain highly sensitive to cytotoxic agents that are not known ABCB1 substrates (exatecan and PNU-159682), both of which have been successfully incorporated as payloads in ADCs.[20,22,23]. These observations raise the possibility that modifying ADC payloads to incorporate non-ABCB1-substrate payloads could circumvent efflux-mediated resistance altogether. Such an approach may be particularly attractive given the potential challenges associated with systemic MDR inhibition, including potential off-target effects and the need for sustained target engagement.

Importantly, our data also indicate that ABCB1-mediated efflux is not the sole mechanism of resistance to VpreB1-ADC. Our ALL1807 cells exhibited relative resistance despite low ABCB1 expression and activity, implicating alternative pathways. *TCF3*::*HLF* B-ALL is also characterized by high expression of anti-apoptotic proteins, particularly BCL2, which may elevate the apoptotic threshold required for ADC-induced cell death[10,12]. Consistent with this, prior work in AML has shown that inhibition of BCL2 can enhance the cytotoxic activity of calicheamicin-based ADCs such as gemtuzumab ozogamicin, supporting a functional role for apoptotic priming in modulating ADC response[43]. In addition, genomic analyses of B-ALL cells resistant to the CD22-directed ADC, inotuzumab ozogamicin, identified diverse, non-efflux-mediated mechanisms that enabled ADC resistance[44,45]. Accordingly, future studies could examine the role of anti-apoptotic programs in VpreB1-ADC resistance and apply unbiased genomic approaches to identify additional mechanisms of resistance.

In summary, this study identifies ABCB1-mediated drug efflux as a key mechanism of VpreB1-ADC insensitivity and demonstrates that this resistance is both reversible and potentially preventable with co-therapeutic modulation. These findings have important implications not only for the clinical development of VpreB1-targeted therapies, but also for the broader field of ADCs in hematologic malignancies. Combining ADCs with MDR inhibitors or employing alternative payloads may represent generalizable strategies to enhance efficacy and overcome resistance, ultimately improving outcomes for patients with high-risk and treatment-refractory B-ALL.

## Acknowledgements

This work utilized the University of Minnesota Masonic Cancer Center shared flow cytometry core which is supported in part by NIH P30 CA77598. We remain grateful to the Simcoe and Simutis families for their support of *TCF3*::*HLF* ALL research studies in honor and memory of Andrew David Simutis.

## Author Contributions

Designed and performed experiments, analyzed data, and prepared figures: Wang X, Ostergaard J, Kang J, Gohman M, Lambert L, Singleton T, Hilgers M, Lee K, Muretta J

Provided critical experimental reagents: Tasian S, Winter S

Designed the study, oversaw the laboratory investigations, performed data analysis, prepared figures, and wrote the manuscript: Williams R, Gordon P

All authors have reviewed, edited, and approved the final version of the manuscript.

## Data Availability Statement

The datasets generated and analyzed during the current study are available from the corresponding author upon request.

## AI Statement

Not applicable.

