## Supplementary material for "ABCB1-Mediated Drug Efflux Drives Resistance to VpreB1-Targeted Antibody-Drug Conjugates in B-cell Lymphoblastic Leukemia": Suppl figures 1-7

**Supplemental Figures and Table**

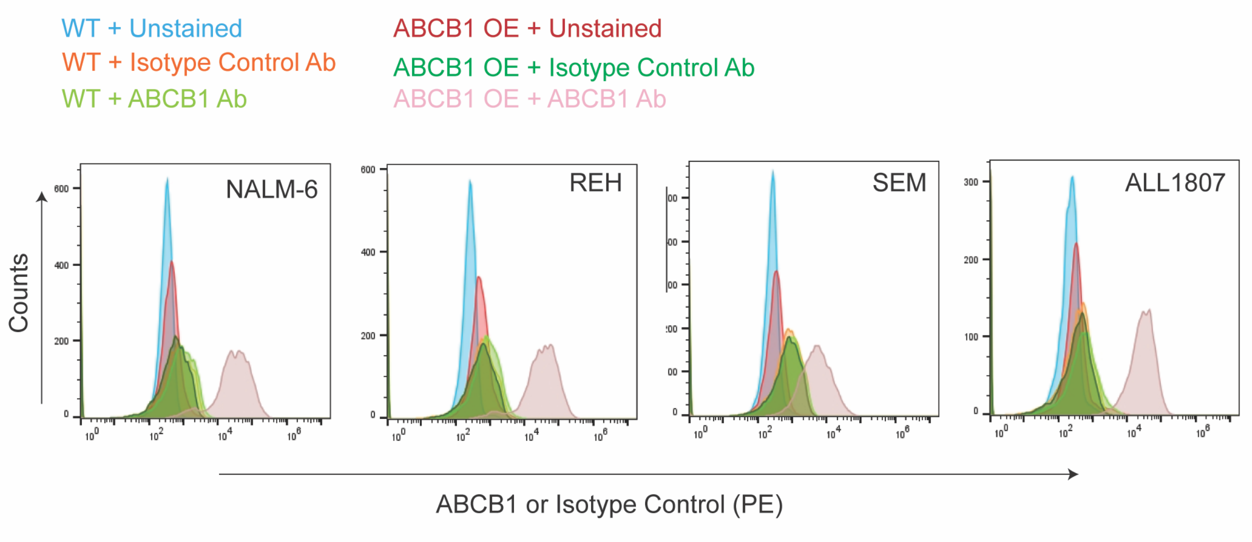


**Supplemental Figure 1:** ***ABCB1 overexpression in leukemia cell lines.*** Leukemia cell lines engineered to overexpress ABCB1 were stained with an anti-ABCB1 antibody or isotype control and analyzed by flow cytometry.

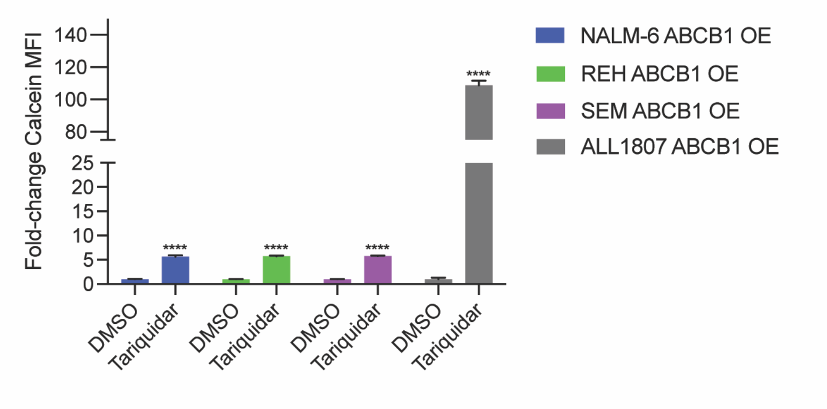


**Supplemental Figure 2:** ***Tariquidar increases calcein accumulation in leukemia cell lines engineered to overexpress ABCB1.*** Median fluorescence intensity (MFI) of intracellular calcein was quantified by flow cytometry in ABCB1-overexpressing leukemia cell lines treated with either DMSO or tariquidar. Error bars represent mean ± SD of three technical replicates. *P* values: ns, not significant; ****, *P* < 0.0001.

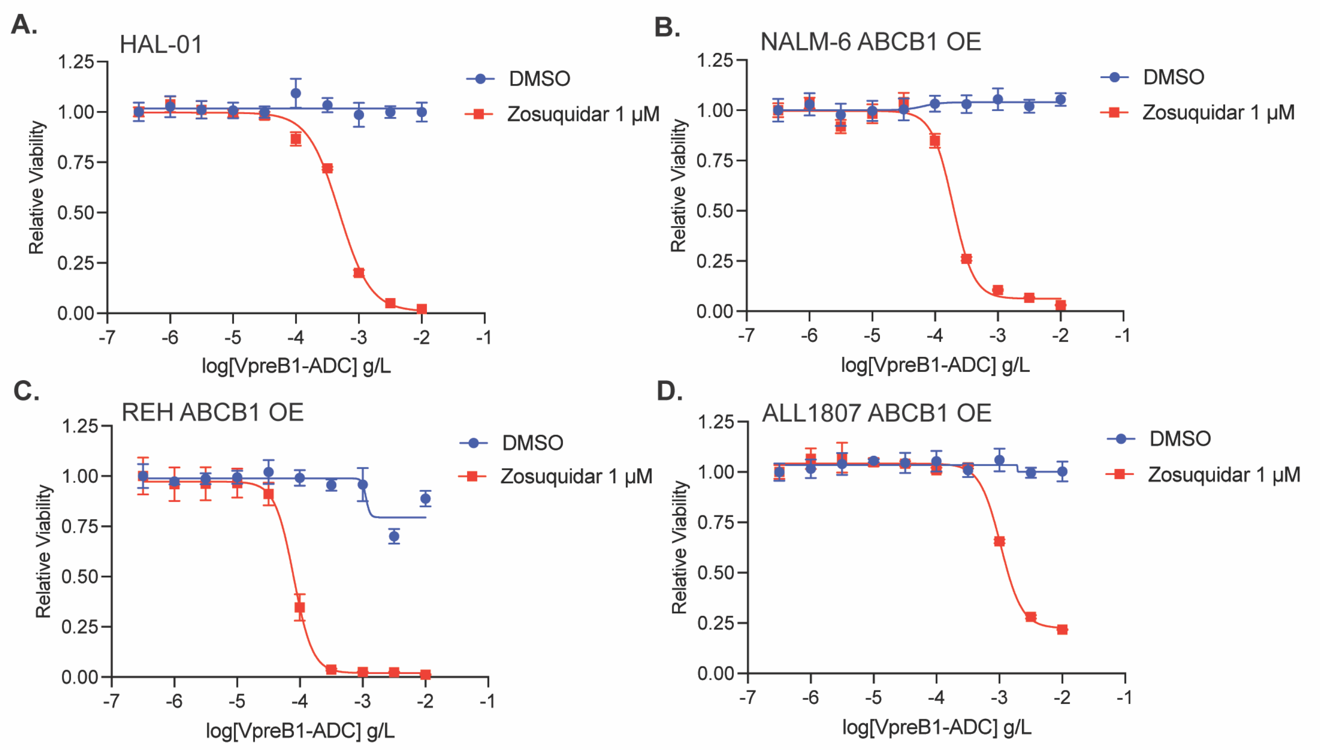


**Supplemental Figure 3:** ***Zosuquidar overcomes ABCB1-mediated VpreB1-ADC resistance***. VpreB1-ADC dose-response curves for HAL-01 (**A**), NALM-6 ABCB1-overexpressing (**B**), REH ABCB1-overexpressing (**C**), and ALL1807 ABCB1-overexpressing (**D**) leukemia cell lines in the presence of zosuquidar 1 μM or DMSO control. Leukemia cell viability was measured after 48 hours using the CellTiter-Glo Assay. Error bars represent mean ± SD of three technical replicates.

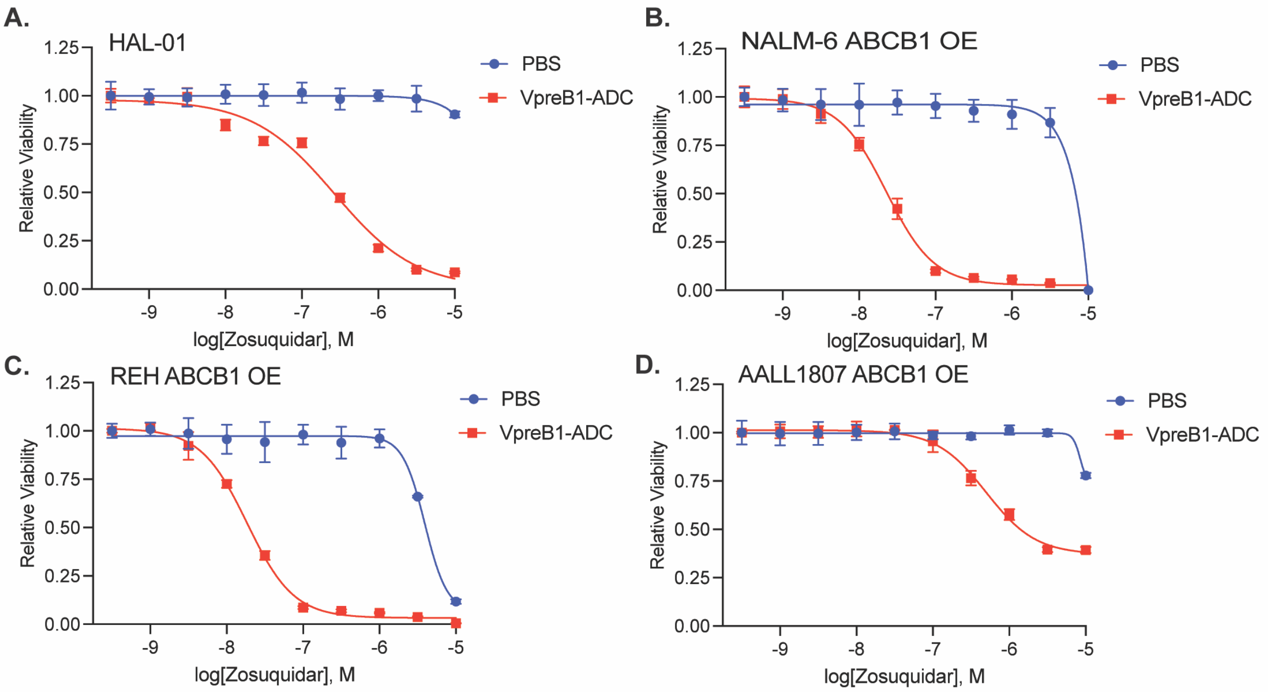


**E.**

| | **log IC50** | **95% CI** |
|---|---|---|
| **HAL-01** | -6.572 | -6.734 to-6.410 |
| **NALM-6 ABCB1 OE** | -7.732 | -7.778 to -7.687 |
| **REH ABCB1 OE** | -7.643 | -7.700 to -7.586 |
| **ALL1807 ABCB1 OE** | -6.303 | -6.389 to -6.217 |

**Supplemental Figure 4: *Zosuquidar $IC_{50}$ determination.* A-D**. Zosuquidar dose-response curves for HAL-01 (**A**), NALM-6 ABCB1-overexpressing (**B**), REH ABCB1-overexpressing (**C**), and ALL1807 ABCB1-overexpressing (**D**) leukemia cell lines in the presence of VpreB1-ADC 0.001g/L or vehicle control. Leukemia cell viability was measured after 48 hours using the CellTiter-Glo Assay. Error bars represent mean ± SD of three technical replicates. **E**. Zosuquidar $IC_{50}$ values calculated from the dose-response curves.

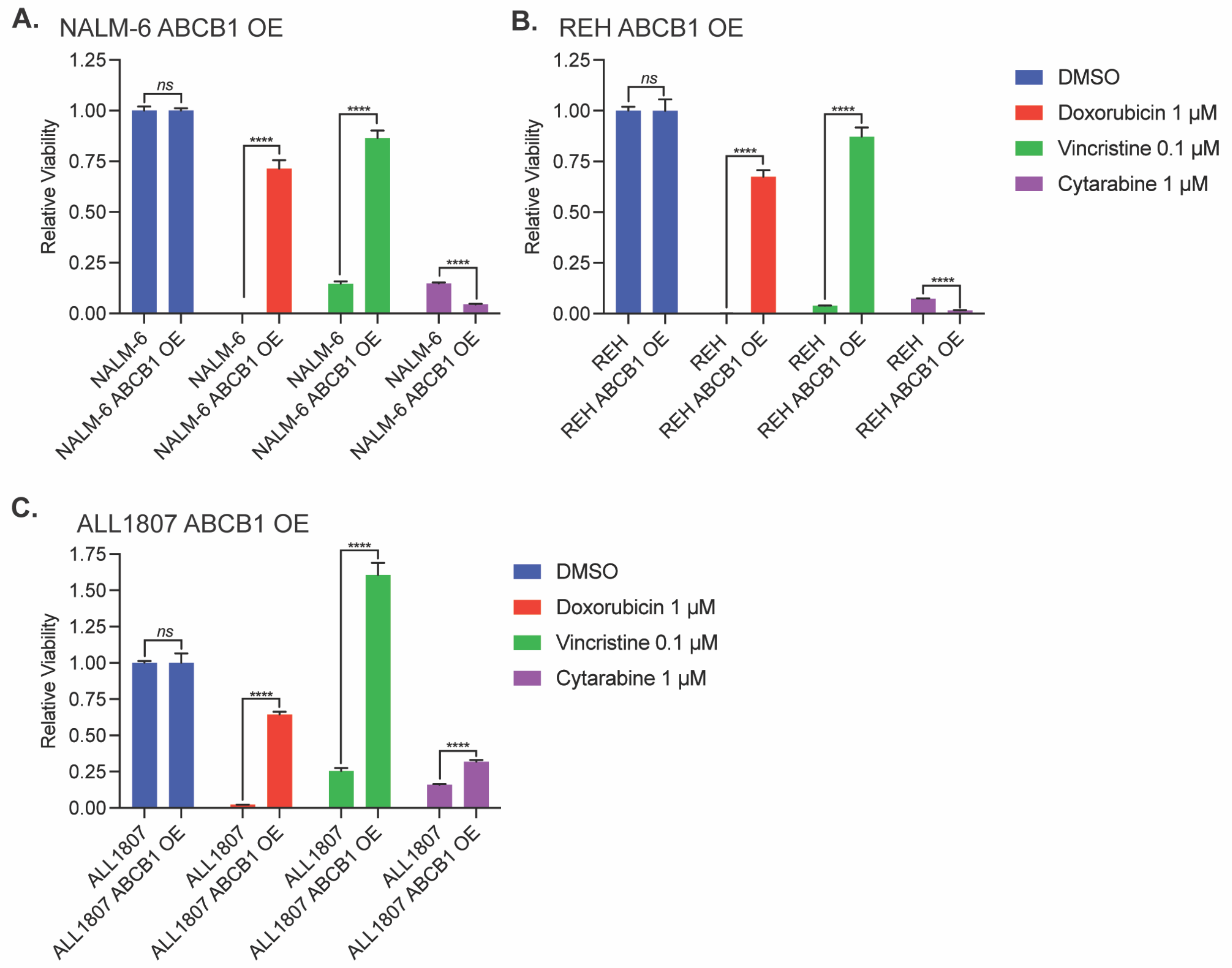


**Supplemental Figure 5:** ***ABCB1 enhances leukemia chemoresistance.*** **A-C**. NALM-6 ABCB1-overexpressing (**A**), REH ABCB1-overexpressing (**B**), and ALL1807 ABCB1-overexpressing (**C**) leukemia cell lines were treated with doxorubicin, vincristine, cytarabine, or DMSO for 48 hours and viability assessed with the CellTiter-Glo Assay. Error bars represent mean ± SD of three technical replicates. *P* values: ns, not significant; ****, *P* < 0.0001.

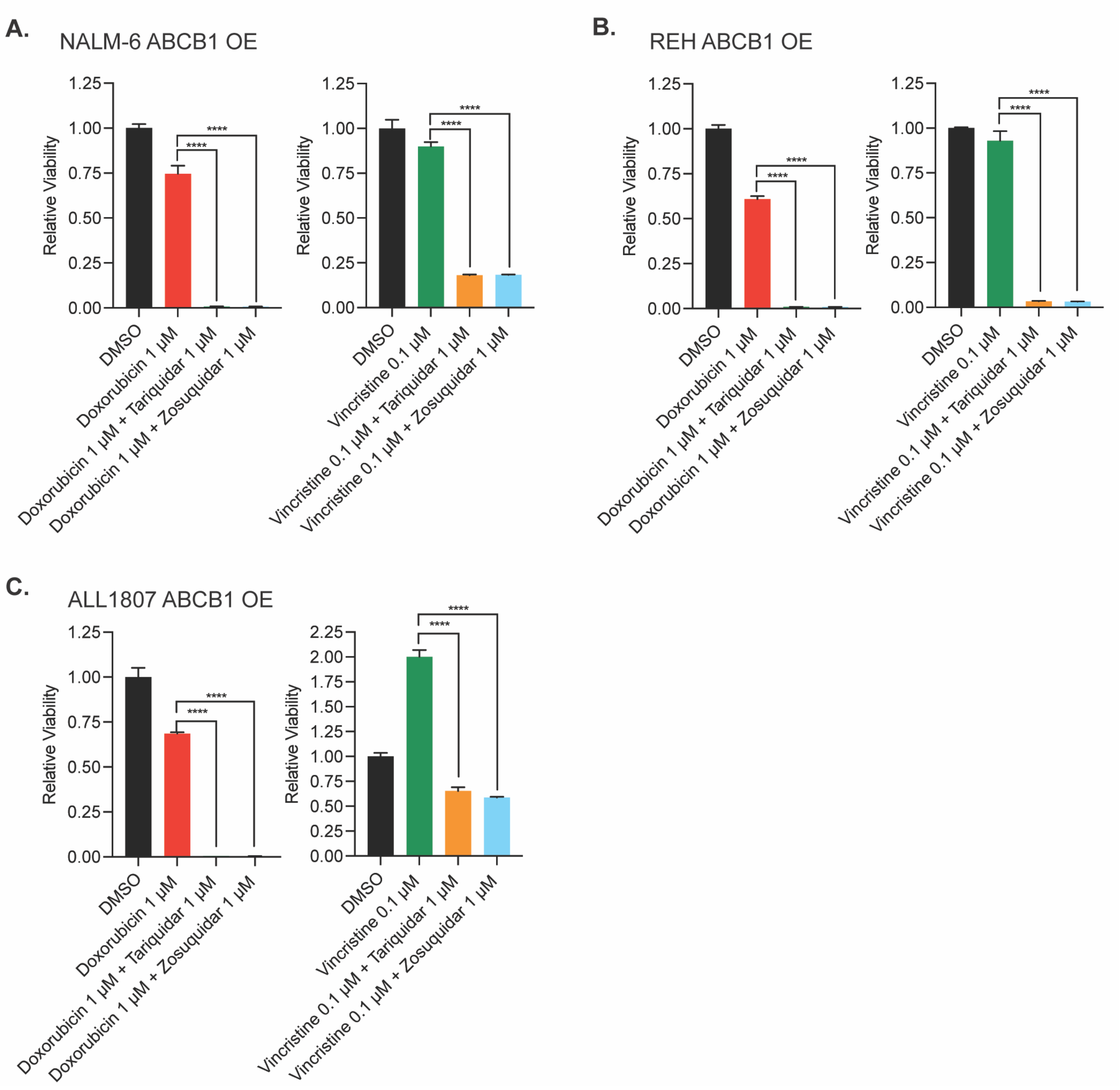


**Supplemental Figure 6**: ***ABCB1 inhibition restores sensitivity to chemotherapy.* A-C**. NALM-6 ABCB1-overexpressing (**A**), REH ABCB1-overexpressing (**B**), and ALL1807 ABCB1-overexpressing (**C**) leukemia cell lines were treated with doxorubicin or vincristine in the presence or absence of the ABCB1 inhibitors tariquidar or zosuquidar for 48 hours, followed by viability assessment using the CellTiter-Glo assay. Error bars represent mean ± SD of three technical replicates. *P* values: ****, *P* < 0.0001.

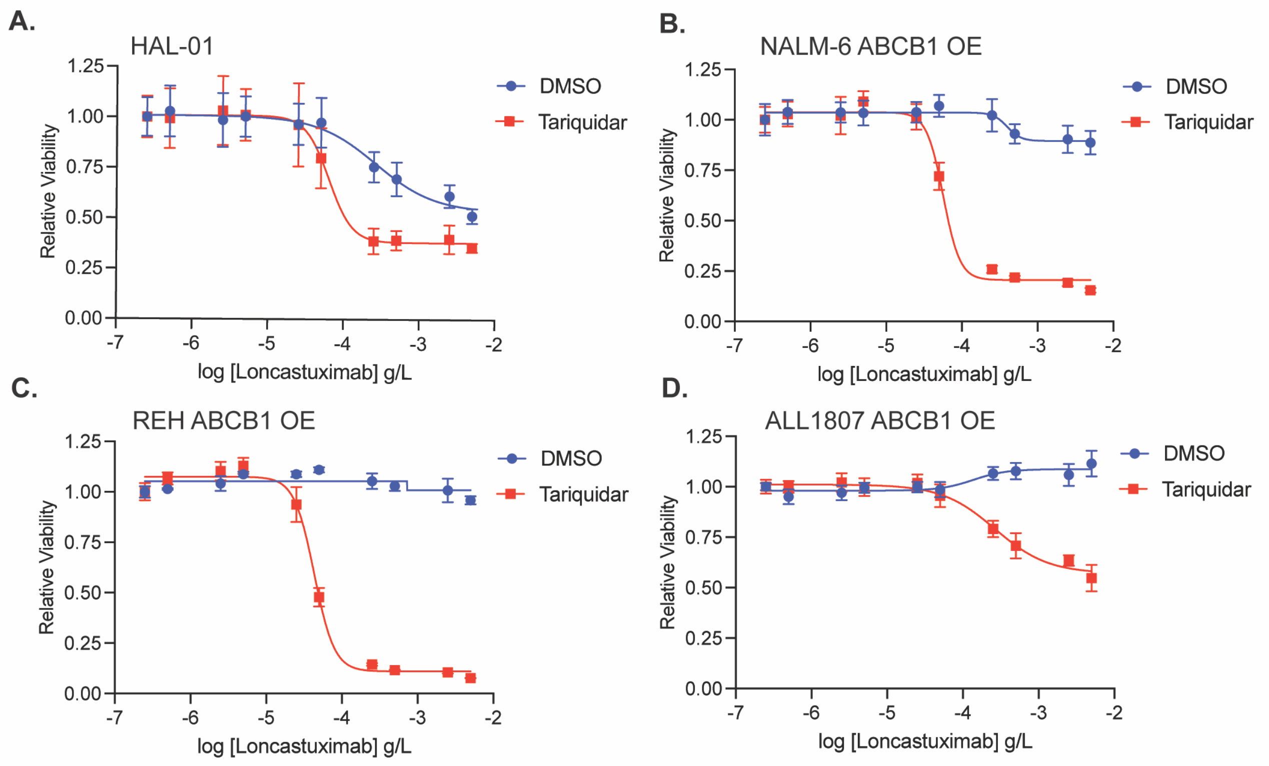


**Supplemental Figure 7:** ***Tariquidar partially overcomes ABCB1-mediated Loncastuximab-tesirine resistance***. Loncastuximab-tesirine dose-response curves for HAL-01 (**A**), NALM-6 ABCB1-overexpressing (**B**), REH ABCB1-overexpressing (**C**), and ALL1807 ABCB1-overexpressing (**D**) leukemia cell lines in the presence of tariquidar 1 μM or DMSO control. Leukemia cell viability was measured after 48 hours using the CellTiter-Glo Assay. Error bars represent mean ± SD of three technical replicates.
